# Macaque-human differences in SARS-CoV-2 Spike antibody response elicited by vaccination or infection

**DOI:** 10.1101/2021.12.01.470697

**Authors:** Alexandra C. Willcox, Kevin Sung, Meghan E. Garrett, Jared G. Galloway, Megan A. O’Connor, Jesse H. Erasmus, Jennifer K. Logue, David W. Hawman, Helen Y. Chu, Kim J. Hasenkrug, Deborah H. Fuller, Frederick A. Matsen, Julie Overbaugh

## Abstract

Macaques are a commonly used model for studying immunity to human viruses, including for studies of SARS-CoV-2 infection and vaccination. However, it is unknown whether macaque antibody responses recapitulate, and thus appropriately model, the response in humans. To answer this question, we employed a phage-based deep mutational scanning approach (Phage-DMS) to compare which linear epitopes are targeted on the SARS-CoV-2 Spike protein in humans and macaques following either vaccination or infection. We also used Phage-DMS to determine antibody escape pathways within each epitope, enabling a granular comparison of antibody binding specificities at the locus level. Overall, we identified some common epitope targets in both macaques and humans, including in the fusion peptide (FP) and stem helix-heptad repeat 2 (SH-H) regions. Differences between groups included a response to epitopes in the N-terminal domain (NTD) and C-terminal domain (CTD) in vaccinated humans but not vaccinated macaques, as well as recognition of a CTD epitope and epitopes flanking the FP in convalescent macaques but not convalescent humans. There was also considerable variability in the escape pathways among individuals within each group. Sera from convalescent macaques showed the least variability in escape overall and converged on a common response with vaccinated humans in the SH-H epitope region, suggesting highly similar antibodies were elicited. Collectively, these findings suggest that the antibody response to SARS-CoV-2 in macaques shares many features with humans, but with substantial differences in the recognition of certain epitopes and considerable individual variability in antibody escape profiles, suggesting a diverse repertoire of antibodies that can respond to major epitopes in both humans and macaques.

**Author summary:** Non-human primates, including macaques, are considered the best animal model for studying infectious diseases that infect humans. Vaccine candidates for SARS-CoV-2 are first tested in macaques to assess immune responses prior to advancing to human trials, and macaques are also used to model the human immune response to SARS-CoV-2 infection. However, there may be differences in how macaque and human antibodies recognize the SARS-CoV-2 entry protein, Spike. Here we characterized the locations on Spike that are recognized by antibodies from vaccinated or infected macaques and humans. We also made mutations to the viral sequence and assessed how these affected antibody binding, enabling a comparison of antibody binding requirements between macaques and humans at a very precise level. We found that macaques and humans share some responses, but also recognize distinct regions of Spike. We also found that in general, antibodies from different individuals had unique responses to viral mutations, regardless of species. These results will yield a better understanding of how macaque data can be used to inform human immunity to SARS-CoV-2.

## Introduction

The COVID-19 pandemic has created a pressing need to understand immunity to SARS-CoV-2, both in the setting of vaccination and infection. This has prompted numerous studies in non-human primates (NHPs), which are considered the most relevant animal model for studying many infectious diseases of humans. Various NHP models have been employed to study the immunogenicity and protective efficacy of SARS-CoV-2 vaccine candidates, with most studies using macaque species including rhesus macaques (*Macaca mulatta*) [1–23], cynomolgus macaques (*Macaca fascicularis*) [8, 24–32], and pigtail macaques (*Macaca nemestrina*) [22, 33–35]. Some of these models have also been used to study infection and re-infection [35–39]. In the NHP model, studies typically measure virus neutralizing antibody responses to vaccination or infection. However, no study has investigated the fine binding specificities of both neutralizing and non-neutralizing SARS-CoV-2 antibodies in macaques and how they compare to the human responses they are meant to model.

Coronaviruses such as SARS-CoV-2 enter host cells using their Spike glycoprotein, which is composed of trimeric S1 and S2 subunits. Receptor-binding S1 homotrimers protrude out from the surface of the virion like a crown, giving this family of viruses its name, while the fusion-mediating S2 trimers anchor the protein to the viral membrane. On S1, the receptor-binding domain (RBD) of SARS-CoV-2 Spike protein binds to angiotensin-converting enzyme 2 (ACE2) on host cells [40, 41]. For subsequent membrane fusion to occur, the Spike protein must be cleaved by host cell proteases at the S1/S2 boundary and at an S2’ site located just upstream of the fusion peptide (FP) of S2 [42], leading to substantial conformational changes that likely unmask new epitopes of S2 to immune cells [43].

Antibodies to SARS-CoV-2 Spike protein are especially interesting as a potential correlate of protection, as they have the capacity to block infection and kill infected cells [44–47]. There has understandably been great interest in studying neutralizing antibodies against the RBD, given that such antibodies can directly block interaction with host cells. While RBD-directed antibodies indeed contribute disproportionately to neutralization [48], the majority of the anti- Spike plasma IgG response in convalescent individuals is directed to epitopes outside of the RBD [49, 50]. RBD-directed antibodies are also less likely to maintain activity against future viral strains, given the increasing number of variants of concern that harbor mutations in the RBD and have reduced sensitivity to neutralization by immune plasma [51]. Additionally, growing evidence from studies in humans and animal models indicates that non-neutralizing antibodies play a role in protection [52–57].

Previous studies have used Phage-DMS [58], a tool that combines phage display of linear epitopes with deep mutational scanning, to interrogate the fine binding specificities and escape profiles of binding antibodies against all domains of Spike in infected and vaccinated humans [59, 60]. These studies have shown that infection-induced human polyclonal antibodies consistently bind linear epitopes in the FP and stem helix-heptad repeat 2 (SH-H) epitope regions, with patient-to-patient variability in escape profiles [59]. Comparatively, mRNA vaccination induces a broader antibody response across Spike protein with more consistent escape profiles [60].

In this study, we built on this foundation by using Phage-DMS to study the binding and escape profiles of antibodies in vaccinated and convalescent macaques in comparison to humans. Our data reveal broad overlap in some major epitopes targeted by both macaques and humans, though neither vaccinated nor convalescent macaques perfectly model the human response. We also find considerable variability in individuals’ antibody escape pathways in most epitope regions in both macaques and humans. The broadest responses were seen in vaccinated humans and re-infected rhesus macaques, groups that also share more concordant escape profiles. These results have implications for the interpretation of COVID-19 macaque research studies.

## Results

Four groups were included in this study: vaccinated pigtail macaques, vaccinated humans, convalescent (re-infected) rhesus macaques, and convalescent humans (Table 1). The vaccinated macaques received a replicating mRNA (repRNA) vaccine encoding the full-length wildtype (not pre-fusion stabilized) SARS-CoV-2 A.1 lineage Spike protein formulated with a cationic nanocarrier [35, 61]. The vaccine was delivered as a prime-only 25ug (n=3) or 250ug (n=6) dose or prime-boost 50ug dose (n=2), with plasma collected 42 days after the first dose (n=9) or 14 days after the second dose (n=2). The vaccinated humans received two doses of the 100ug Moderna mRNA-1273 vaccine encoding the pre-fusion stabilized full-length SARS-CoV-2

**Table 1.** Details of samples used in the current study.

| Group | Number of samples | Age range (years) | Treatment | Time of sample collection |
| --- | --- | --- | --- | --- |
| Vaccinated pigtail macaques | 11 | 3 ½ - 6 | repRNA vaccine encoding full-length SARS-CoV-2 Spike<sup>a</sup> | 42 days post 1<sup>st</sup> dose |
| Vaccinated humans | 15 | 18 - 55 | 100ug mRNA vaccine encoding full-length pre-fusion stabilized SARS-CoV-2 Spike (Moderna) | 36 days post 1<sup>st</sup> dose |
| Convalescent rhesus macaques | 12 | 2 ½ - 5 | Infected twice with SARS-CoV-2 six weeks apart<sup>a</sup> | 56 days post 1<sup>st</sup> infection |
| Convalescent humans | 12 | 28 - 52 | Naturally infected once with SARS-CoV-2 (mild disease) | Median 67 (IQR 62, 70) days post symptom onset |

<sup>a</sup>Within each group of macaques, subgroups received slightly different treatments (described in Table S1).

A.1 lineage Spike protein and formulated with a lipid nanoparticle. Serum was collected from human vaccinees 36 days after the first dose (7 days after the second dose). The convalescent macaques were infected twice with SARS-CoV-2, with infections spaced six weeks apart and serum collected 56 days after the first infection (14 days after the second infection). The convalescent humans were naturally infected once with SARS-CoV-2 and exhibited mild disease, with a median of 67 days between symptom onset and sample collection. Details of individual participants are available in Table S1.

### Enrichment of wildtype peptides

To compare which regions of Spike protein are recognized by human and macaque antibodies, we examined the enrichment of wildtype peptides by antibodies from each individual (Fig 1A). Broadly speaking, binding was observed in the NTD, CTD, FP, and stem helix-HR2 epitope regions as reported previously in human studies [59, 60]. Epitope regions (shown as different colors on Fig 1) were defined as previously [60]: NTD, amino acid 285-305; FP, 805-835; stem helix-HR2 (SH-H), 1135-1170. For the CTD, the bounds of epitope regions were expanded and altered from previous studies based on macaque antibodies recognizing a wider area than previously seen in humans: CTD-N’, 526-593; CTD-C’, 594-685 (S1A Fig). Several additional epitopes that flank previously-defined regions were also identified in this analysis: pre-FP, 777- 804; post-FP, 836-855 (S1B Fig); and HR2, 1171-1204 (S1C Fig). Specific epitope regions can be visualized on the structure of a Spike protein monomer in Fig 1B. In addition to these defined regions, we noted that one convalescent rhesus macaque appeared to weakly recognize an epitope at the beginning of the S2 subunit (amino acid 686-710, Fig 1A).

**Fig 1:**
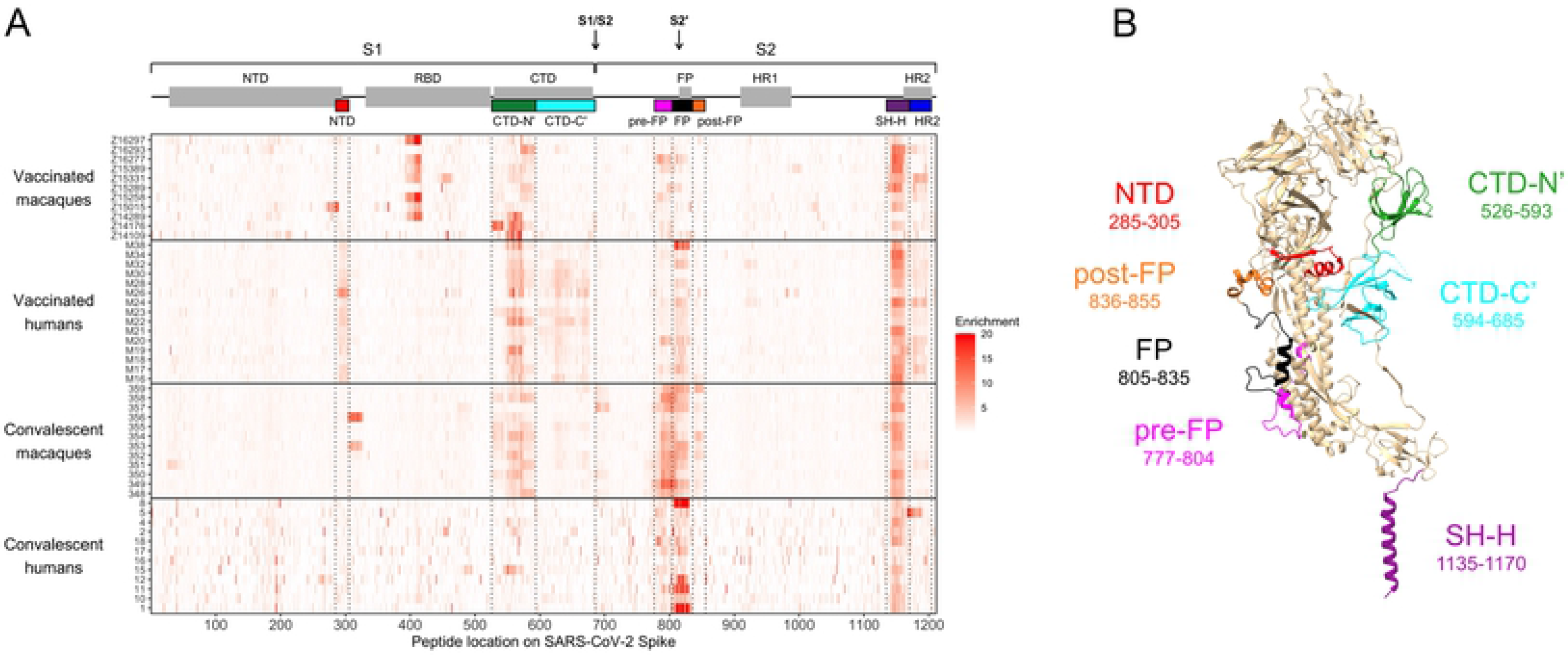
Enrichment of wildtype peptides. (A) The x axis indicates each peptide’s location along SARS-CoV-2 Spike protein, and each entry on the y axis is an individual sample. All enrichment values over 20 are plotted as 20 to better show the lower range of the data. Above the heatmap, domains of Spike are shown with grey boxes, with the S1/S2 and S2’ cleavage sites indicated with arrows. The epitope regions defined in the current study are shown as colored boxes (from left to right: NTD in red, CTD-N’ in green, CTD-C’ in cyan, pre-FP in pink, FP in black, post-FP in orange, SH-H in purple, and HR2 in blue). (B) Defined epitope regions shown on a structure of one monomer of SARS-CoV-2 Spike in the pre-fusion conformation (PDB 6XR8 [ref 95]). The amino acid loci spanned by each epitope are listed. The HR2 epitope (AA 1171-1204) could not be resolved on the structure and is not shown.

In general, we did not detect responses in the RBD because many epitopes in this region are known to be conformational, and Phage-DMS only has the power to detect epitopes that include linear sequences. Epitopes in the RBD have been extensively detailed elsewhere [62, 63]. However, we did detect strong binding to an RBD epitope in some vaccinated pigtail macaques (Fig 1A). This same region was enriched in samples from before vaccination in four of the five pigtail macaques with baseline samples available (S2 Fig). Pre-infection serum from the twelve rhesus macaques did not show any consistent responses (S2 Fig). Because the RBD response in pigtail macaques was present prior to vaccination with SARS-CoV-2 Spike, we did not investigate it further as a response to vaccination.

To quantify differences in the epitopes targeted by different groups, the enrichment of wildtype peptides was summed across each epitope region for every individual. Because the main research question is whether responses in macaques model those in humans, two comparisons were performed: vaccinated macaques vs. vaccinated humans and convalescent macaques vs. convalescent humans (Fig 2).

**Fig 2:**
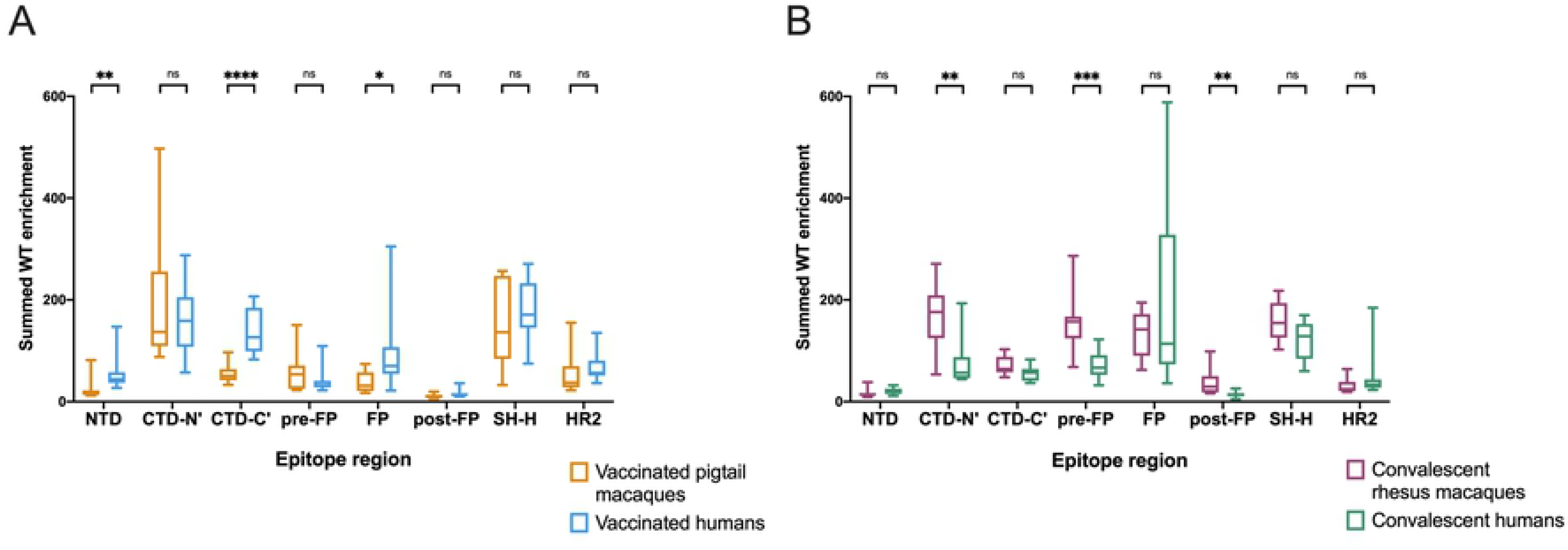
Differences in enrichment of wildtype peptides by group. Wildtype enrichment values were summed for all peptides within each epitope region. Box plots summarize the data by group. (A) compares vaccinated pigtail macaques to vaccinated humans, while (B) compares convalescent rhesus macaques to convalescent humans. Multiple Mann-Whitney U tests were performed, with p values corrected for the number of comparisons in each plot (8) using the Bonferroni-Dunn method. ****, p ≤ 0.0001; ***, p ≤ 0.001; **, p ≤ 0.01; *, p ≤ 0.05.

In concordance with a qualitative assessment of the enrichment heatmap in Fig 1A, vaccinated humans preferentially recognized the following epitope regions compared to vaccinated macaques: NTD (Mann-Whitney p ≤ 0.01), CTD-C’ (p ≤ 0.0001), and FP (p ≤ 0.05) (Fig 2A).

Meanwhile, convalescent macaques recognized the following epitope regions more than convalescent humans: CTD-N’ (p ≤ 0.01), pre-FP (p ≤ 0.001), and post-FP (p ≤ 0.01) (Fig 2B). All groups consistently recognized the SH-H epitope region (Fig 2). While vaccination appeared to induce a stronger response against HR2 than infection (Fig 1A), there were no significant differences in response driven by species (Fig 2). Within each group of macaques (vaccinated and convalescent), subgroups received slightly different treatments (Table S1), so similar analyses were performed comparing these subgroups; no comparisons were significant at a threshold of p=0.05 (Kruskal-Wallis test, S3 Fig).

Taken together, these findings indicate: 1) vaccinated humans were the only group to consistently recognize peptides from both the NTD and CTD-C’ epitope regions, which are in close physical proximity to one another (Fig 1B); 2) convalescent humans had a limited response to the CTD-N’; 3) compared to other groups, convalescent macaques had a notably more robust response to regions upstream and downstream of the main FP epitope region; 4) vaccinated macaques did not recognize the FP as strongly as other groups; and 5) vaccination seemed to induce a stronger response against HR2 than infection in both macaques and humans.

### Defining and comparing escape pathways

To assess differences in the binding characteristics of human and macaque antibodies on a more granular level, we next examined the mutations in Spike that reduced antibody binding in each epitope region of interest. Because the antibody escape pathways for vaccinated humans have been described previously [60], we did not examine the NTD and CTD-C’, which are exclusively recognized by this group. Instead, we focused on comparing escape profiles between groups in the following epitope regions: CTD-N’, FP, and SH-H. We first represent the data as scaled differential selection values in logo plot form, as commonly shown in previous studies. Importantly, scaled differential selection is highly correlated with peptide binding as measured by competition ELISA [58]. To summarize the data represented by the logo plots by group, summed differential selection values across each epitope region were also calculated. This metric represents the overall magnitude of escape at each locus regardless of the specific amino acid substitution, with negative values indicating a decrease in binding compared to the wildtype amino acid, and positive values indicating enhanced binding (see “Materials and Methods”). Finally, escape similarity scores were calculated between pairs of individuals to quantify similarity in escape profiles (see “Materials and Methods” and S4 Fig).

### CTD-N’

Vaccinated macaques, vaccinated humans, and convalescent macaques recognized peptides in the CTD-N’ (AA 526-593), whereas convalescent humans generally did not (Fig 2B). Within this epitope region, the individual escape profiles showed notable variability both within and between groups (S5 Fig). For example, across all groups, some individuals showed relatively high sensitivity to mutations between sites 558-567, while others had a response focused more downstream around AA 577-586. There was also substantial variability in which loci in the CTD-N’ had the highest relative magnitude of escape, and sometimes even in the directionality of scaled differential selection at a given locus. For example, some individuals had antibodies that bound mutated peptides better than wildtype at AA 555 (e.g., convalescent macaque 353) while others exhibited reduced binding to mutated peptides (e.g., convalescent macaque 358). The same was true for site 560 (e.g., vaccinated humans M24 and M26 exhibited improved and disrupted binding to mutated peptides, respectively).

To summarize the trends observed in the individual findings, we calculated summed differential selection values for each individual at each site and generated boxplots by group (Fig 3A). In addition to the aforementioned regions of escape common to all groups, convalescent macaques also showed considerable escape between AA 529-535, with vaccinated macaques also showing a less consistent response in this area (Figs. 3A and S5). The complexity and variability of the escape pathways also prompted us to quantify the similarity in escape between and within groups. Escape similarity scores largely corresponded to areas of high magnitude of escape. Sites with low-magnitude summed differential selection values indicate loci where mutations have no notable impact, and therefore those escape profiles reflect fluctuations in peptide enrichments due to noise, which drives a lower escape similarity score at those sites (Fig 3A, lower panel). At some sites (e.g., 560, as described above), low scores were also the result of some samples showing negative differential selection and others showing positive differential selection, a comparison that was assigned the highest cost in our escape similarity score algorithm.

**Fig 3:**
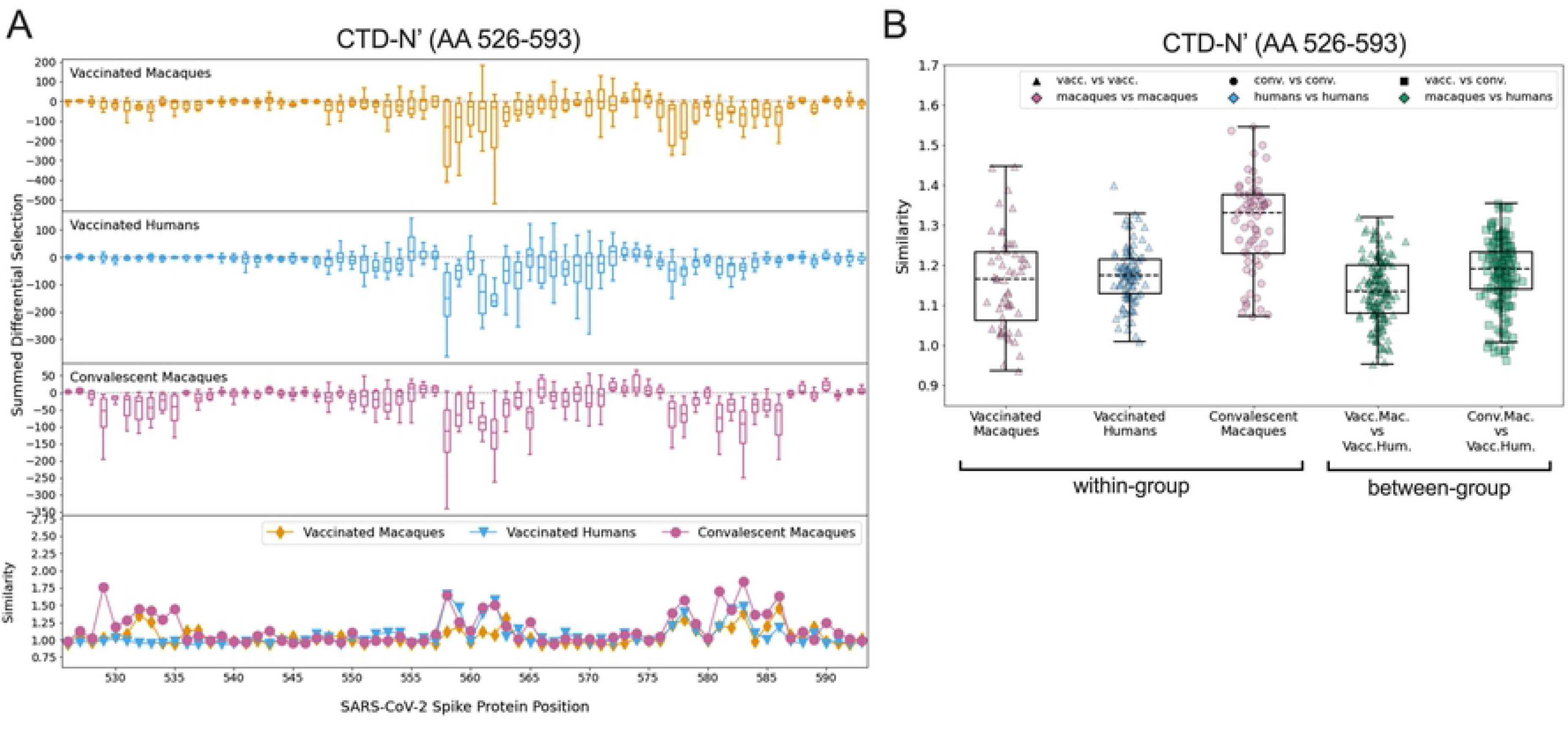
Comparison of escape profiles in the CTD-N’. (A) The top three panels show boxplots depicting the summed differential selection values of all samples in a group at each locus. Negative values represent sites where the binding interaction between antibody and peptide was weakened when peptides were mutated, whereas positive values represent enhanced binding. The bottom panel shows the mean escape similarity score for all pairwise comparisons between samples in each group, calculated at every locus. See S4 Fig for a description of the escape similarity score algorithm. (B) Within- and between-group region-wide escape similarity scores, summarized as boxplots. Each point represents a pairwise comparison between two samples. The contribution of a site’s score to the total escape similarity score is weighted based on its relative contribution to the summed differential selection values across the region. P values are not computed due to lack of independence between data points.

To test the similarity of escape profiles across the CTD-N’ epitope region, escape similarity scores were aggregated across the region and computed both within and between groups. These are shown as boxplots, with each point representing a pairwise comparison between individual samples (Fig 3B). For example, every vaccinated macaque was compared to every other vaccinated macaque (a within-group comparison) and to every vaccinated human (a between-group comparison). We included a comparison of convalescent macaques and vaccinated humans, given visual similarities between their patterns of escape (Fig 3A).

Convalescent macaques showed the highest within-group similarity in escape profiles, meaning their escape profiles were more consistent than those of the vaccinated macaques or vaccinated humans (Fig 3B). Between-group escape similarity scores were on par with the within-group scores for the vaccinated macaques and humans, indicating that although there was substantial variability in individual profiles, this was not driven by sample groups.

### FP

Escape profiles were examined in the FP epitope region (AA 805-835) for the three groups that showed significant wildtype enrichment in this area: vaccinated humans, convalescent macaques, and convalescent humans. As in the CTD-N’, overall there was variability in individual escape profiles, though the convalescent macaques showed a more consistent pattern of escape than other groups (S6 Fig). Within the FP, most sites of escape fell between AA 811-825 for all groups (Fig 4A). The convalescent macaques again exhibited the highest escape similarity scores (Fig 4B). The median within-group escape similarity scores in the FP were on par with those in the CTD-N’ (Fig 3B), indicating approximately equal variability in antibody escape in these epitope regions. The between-group escape similarity scores were generally similar to each other and to the human within-group scores (Fig 4B).

**Fig 4:**
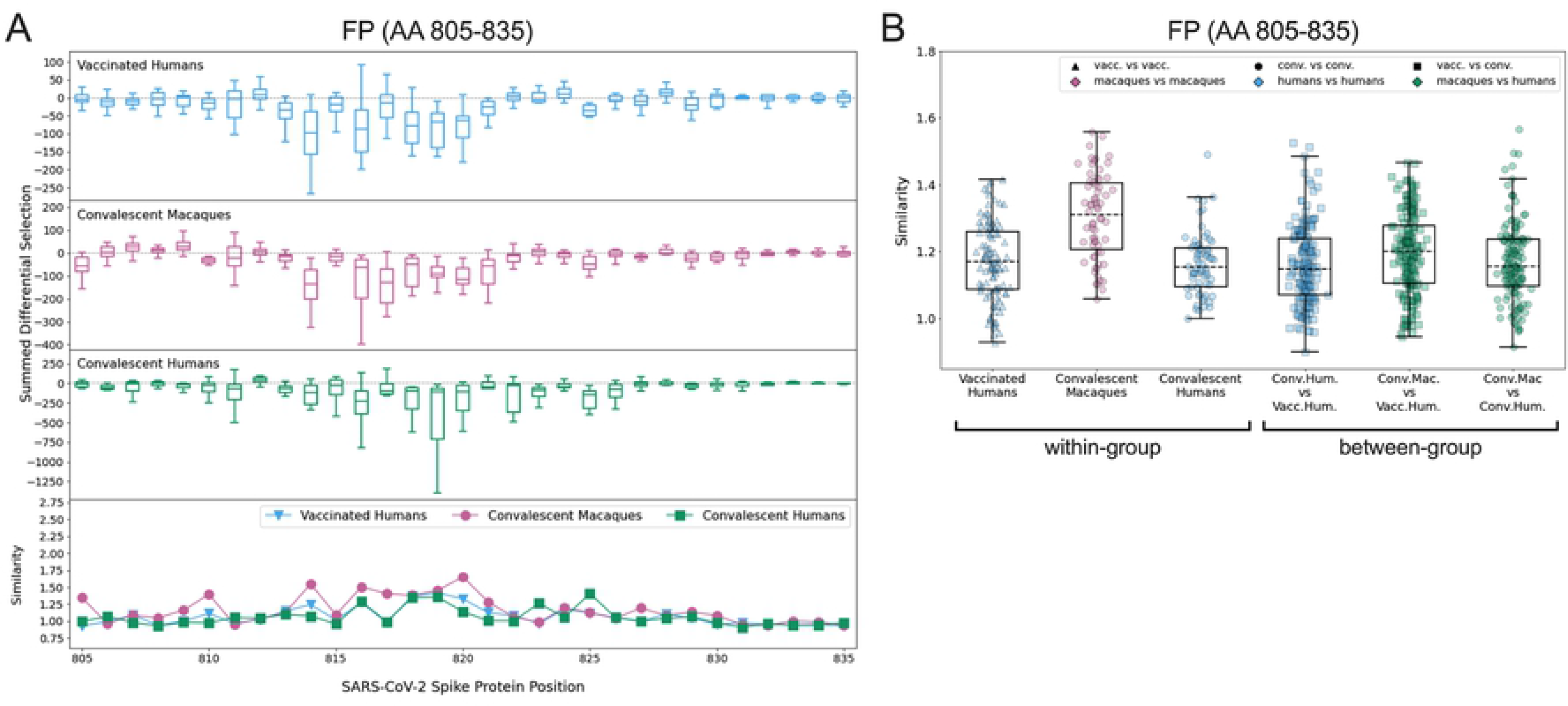
Comparison of escape profiles in the fusion peptide (FP). (A) and (B) Data are shown as described in Fig 3.

### SH-H

All four groups consistently recognized peptides spanning the SH-H epitope region (AA 1135- 1170). Major sites of escape were located between AA 1145-1158 for all groups (Fig 5A). The individual logo plots in the SH-H suggested a consistent response among vaccinated humans and convalescent macaques, with more variability in the remaining groups (S7 Fig). This finding is supported by the within-group escape similarity scores for those groups trending higher across the epitope region (Figs. 5A lower panel and 5B). The median epitope region-wide escape similarity scores for vaccinated humans and convalescent macaques were also higher in the SH-H than in the CTD-N’ or FP, confirming a more concordant response. The median between-group escape similarity score for vaccinated humans and convalescent macaques was on par with their median within-group scores, indicating that the escape profile of a vaccinated human looks as similar to that of a convalescent macaque as it does to another vaccinated human (Fig 5B). The similarity between these two groups was higher than the similarity between convalescent macaques and humans, as well as between vaccinated macaques and humans (Fig 5B). Despite this overall trend, two vaccinated humans had more unique escape profiles (S7 Fig, M26 and M19) and are responsible for a cluster of lower-similarity outlier points (Fig 5B, “Vaccinated Humans” and “Conv. Mac. vs. Vacc. Hum.”).

**Fig 5:**
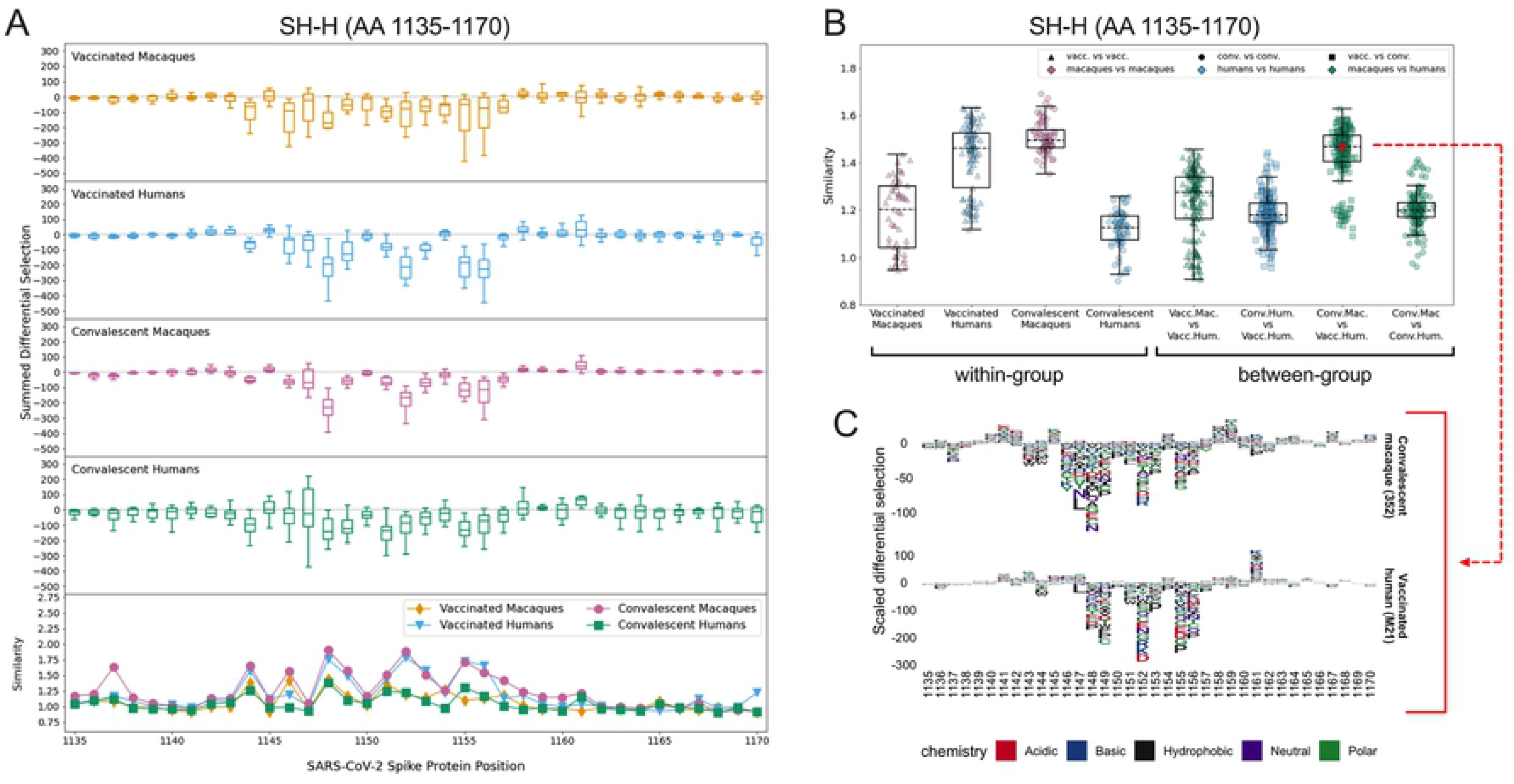
Comparison of escape profiles in the stem helix-HR2 region (SH-H). (A) and (B) Data are shown as described in Fig 3. (C) Logo plots for participant 352 (a convalescent macaque) and M21 (a vaccinated human) showing the effect of specific mutations on antibody binding at each site. The comparison between these samples had an escape similarity score closest to the median value for all pairwise convalescent macaque vs. vaccinated human comparisons and thus can be considered representative of the similarity between these groups. The 352 – M21 comparison is shown in red on (B).

The pairwise comparison between participant 352 (a convalescent macaque) and M21 (a vaccinated human) generated an escape similarity score closest to the median for all comparisons between these groups. Logo plots for these individuals are shown in Fig 5C as a representative example of the striking between-group similarity. The most consistent sites of escape for both groups were AAs 1148, 1152, 1155, and 1156 (Figs. 5A and S7). While some differences exist, there was not nearly as much variability as in the CTD-N’ (S5 Fig) and FP (S6 Fig).

### Other epitope regions

In addition to the epitope regions described above, the convalescent macaques strongly recognized the pre-FP and post-FP, which were not targeted by human antibody responses (S8 Fig). Escape profiles in the pre-FP appeared highly consistent among individual macaques, with major sites of escape at AAs 795, 798, 800, and 802. Profiles were more variable in the post-FP, likely due in part to low enrichment of wildtype peptides in this epitope region for some individuals (S8 Fig).

### Comparison of vaccinated humans and convalescent macaques

It was notable that the vaccinated humans and convalescent macaques showed the most similarity in escape profiles across all epitope regions, most strikingly in the SH-H. Thus, we also asked whether they showed similarity in the epitopes they targeted by comparing the enrichment of wildtype peptides in these groups in each epitope region (S9 Fig). Vaccinated humans recognized the following epitope regions more strongly than convalescent macaques: NTD (Mann-Whitney p ≤ 0.0001), CTD-C’ (p ≤ 0.0001), and HR2 (p ≤ 0.001). Convalescent macaques preferentially recognized the pre-FP (p ≤ 0.0001) and post-FP (p ≤ 0.001) epitope regions. This suggests some diversity in the epitopes targeted, but similarity of antibody escape patterns within epitopes targeted by both groups.

## Discussion

In this study, we aimed to assess whether the antibody binding specificities to SARS-CoV-2 Spike in macaques are a useful model for the human response. Our results indicate important similarities between macaques and humans; for example, both have antibodies that recognize major epitopes in the CTD, FP, and SH-H. However, many differences are also apparent, with some groups showing responses to unique epitopes, such as two physically proximal epitopes in the NTD and CTD that are recognized by antibodies from vaccinated humans but not macaques. Additionally, epitope regions flanking the FP were recognized by antibodies from convalescent macaques, while antibodies from convalescent humans did not recognize the flanking regions but showed a strong response within the FP itself. We found considerable diversity in the pathways of escape between individuals, and this was not specific to either macaques or humans, suggesting a diverse repertoire of antibodies that can respond to the major epitopes in both groups. Overall, these results suggest that macaques and humans share recognition of certain major epitopes. The differences that exist could be due to species (macaque vs. human), but could also be influenced by differences in the specific type and number of exposures to antigen in each group.

Other studies have characterized human monoclonal antibodies against some of the epitopes we report here, many of them with neutralizing or other activities. As previously reported by our group [60], we found that antibodies from vaccinated humans bound peptides spanning a 30 amino acid segment at the C-terminus of the NTD. Interestingly, most if not all neutralizing human mAbs targeting the SARS-CoV-2 NTD to date have been shown to target a single supersite on the “tip” of Spike, distinct from the epitope we detected at the C-terminus [49, 64–70]. An NTD mAb with Fc effector function [71], as well as several NTD mAbs that enhance infection in vitro [65, 72], also bind sites upstream of the C-terminal epitope. Therefore, future studies are warranted to investigate the function of antibodies binding the new NTD epitope detected by Phage-DMS. In the CTD, we detected broad antibody binding, with vaccinated macaques, vaccinated humans, and convalescent macaques enriching peptides in the CTD-N’ epitope region, and vaccinated humans also recognizing peptides spanning the remainder of this domain (CTD-C’). Polyclonal antibodies targeting sites within the CTD-N’ and CTD-C’ have been isolated from human sera and shown to have neutralizing activity [73]. Interestingly, the neutralizing epitope on the CTD-C’ (AA 625-636) [73] is physically adjacent to the NTD epitope we describe (AA 285-305), raising the possibility that a conformational epitope extending to the NTD is recognized by neutralizing antibodies from vaccinated humans. Depleting human serum of FP-binding antibodies reduced its neutralization capacity [74]; these antibodies are of high interest, both due to their potential to block membrane fusion, and given the high sequence conservation among the FPs of diverse coronaviruses [75, 76]. We found that convalescent rhesus macaque sera strongly recognized the pre- and post-FP epitope regions, but to our knowledge, functional antibodies against these regions have not been previously described.

Finally, the SH-H epitope region we describe is in the stem helix, a region known to be highly conserved across coronaviruses. Broadly neutralizing [77–79] stem helix antibodies have been isolated and suggest an avenue for rational design of a pan-coronavirus vaccine. Interestingly, a mAb raised against the MERS-CoV stem region protected mice against SARS-CoV-2 challenge, despite having no neutralizing activity against SARS-CoV-2 in vitro [80]. The detection of broad antibody binding across Spike supports the continued investigation of non-RBD epitopes, which remain understudied. Some of the epitopes we describe may also be the target of non-neutralizing Fc-effector antibodies [81], and/or antibodies that enhance infection via Fc-independent [72] or Fc-dependent [82] mechanisms. This latter concept may be important in the pathogenesis of COVID-19, though this remains speculative.

Previous work elucidated that pathways of antibody escape to SARS-CoV-2 Spike protein can be quite variable in convalescent humans, with vaccination inducing a more consistent response [60]. In the current study, we found considerable variability in escape profiles in the FP and CTD-N’ in both macaques and humans, though the convalescent rhesus macaques had more concordant escape profiles than other groups. Variability in escape patterns suggests that a diversity of antibodies are targeting these epitopes. Intra-species germline diversity in immunoglobulin genes may help explain why individuals with similar exposures often mount distinct responses [83, 84]. On the other hand, escape profiles were more consistent in the SH- H, where the responses of convalescent macaques and vaccinated humans appeared to converge. This conservation of response suggests that highly similar antibodies are dominating the antibody repertoire against this epitope. Convergent antibody responses to SARS-CoV-2 have been reported within human populations [85–87], and our findings here suggest that antibodies from different species may also be able to converge on the same “public” antibody repertoires in a functional sense, despite genetic differences. While a shared escape profile among individuals could suggest that viral escape mutations are more likely to emerge on a population level, another factor to consider is the effect of the mutations on viral fitness. Key domains of the S2 subunit (such as the SH-H epitope) have essential functions and high sequence conservation, suggesting a low tolerance for mutation and thus for escape. Indeed, previous work determined that sites of escape identified by Phage-DMS are not typically mutated at a high frequency in circulating strains of SARS-CoV-2 [59].

While our focus was on understanding how macaques and humans respond to a similar exposure (i.e., vaccination or infection), we also noted similarities in response between re- infected macaques and vaccinated humans. These groups both exhibited the broadest recognition across Spike, although the epitope regions they targeted were somewhat different. As described above, these groups also had highly similar antibody escape profiles in the SH-H. The vaccinated humans and re-infected macaques both received two exposures to high doses of antigen. It is plausible that re-exposure directed initially diverse antibodies to converge on a more focused response in both scenarios. While it is known that vaccination and infection induce distinct humoral responses against Spike [60, 88, 89], our data suggest that a second exposure may generate antibodies that better match the vaccine-induced response.

This study had several limitations. Because the Phage-DMS library displays peptides 31AA in length, discontinuous or conformational epitopes are not readily detected using this method. Additionally, epitopes that may normally be glycosylated are exposed for antibody binding in Phage-DMS. There also are known germline-encoded differences in the properties of immunoglobulin subclasses and Fc receptors between macaques and humans, leading to differences in antibody function that cannot be assayed using Phage-DMS [90]. Additionally, our sample set includes variables that limit our ability to draw conclusions about species- specific (macaque vs. human) differences in antibody response. The vaccinated macaques and humans both received RNA vaccines encoding full-length Spike protein, but there were differences in vaccine technology, including: 1) the use of mRNA in the human vaccine vs. repRNA in the macaque vaccine, 2) the stabilization of Spike in its pre-fusion state in the human vaccine, 3) the dosage and number of doses delivered, and 4) the formulation used to deliver the RNA. Despite these differences, we found commonalities in some of the epitopes targeted by antibodies from both groups. Additionally, the convalescent rhesus macaques were experimentally infected twice with high titers of virus, compared to the convalescent humans who were naturally infected once. This important discrepancy could be the reason why the response in re-infected macaques aligned more closely with vaccinated humans than convalescent humans. Studies of re-infected humans would help address this possibility.

Our findings suggest that while vaccinated and convalescent macaques and humans share recognition of some major epitopes, each group has a unique antibody binding profile.

Antibody escape profiles suggest a diversity of individual responses to most epitopes. Important avenues for future study include comparing macaque and human responses to the RBD and evaluating species differences in antibody function. Continued investigation of immunogenic epitopes in conserved regions of Spike is also warranted to inform the development of immunity that is more robust in the face of viral escape.

## Materials and Methods

### Samples

#### Vaccinated pigtail macaques

Plasma was collected from 11 pigtail macaques immunized with a replicating RNA (repRNA) vaccine expressing full-length SARS-CoV-2 Spike protein. A subset of these animals was previously described [35]. All animals were housed at the Washington National Primate Research Center (WaNPRC), an accredited facility of the American Association for the Accreditation of Laboratory Animal Care International (AAALAC). All procedures were approved by the University of Washington’s Institutional Animal Care and Use Committee (IACUC) (IACUC #4266-14). Individual macaques received the vaccine by intramuscular injection in either a Lipid InOrganic Nanoparticle (LION) [35] or a Nanostructured Lipid Carrier (NLC) [61] formulation, delivered in a single priming dose of 25ug (n=3) or 250ug (n=6) or in a prime-boost regimen with 50ug doses spaced 4 weeks apart (n=2). All samples were collected 6 weeks post-prime immunization. A subset of these animals also previously received an experimental hepatitis B vaccine as part of another study (n=5).

#### Convalescent rhesus macaques

Serum was collected from 12 rhesus macaques housed at the Rocky Mountain Laboratories (National Institutes of Health [NIH]), 14 days after the second of two SARS-CoV-2 infections spaced 42 days apart. Prior to infection, macaques were variably depleted of CD4+ T cells, CD8+ T cells, CD4+ and CD8+ T cells, or neither, as part of another study. Details of macaque treatment and regulatory approvals are as published previously [39].

#### Vaccinated humans

We obtained serum from 15 individuals who received two 100ug doses of the Moderna mRNA- 1273 vaccine as part of a phase I clinical trial (NCT04283461) [91]. Phage-DMS results from these samples were reported previously [60]. Because samples were de-identified, this study was approved by the Fred Hutchinson Cancer Research Center Institutional Review Board as nonhuman subjects research. Only samples from individuals aged 18-55 years were included in the current study to better match the young age range of the macaques.

#### Convalescent humans

Plasma was collected from 12 individuals post-mild COVID-19 illness as part of the Hospitalized or Ambulatory Adults with Respiratory Viral Infections (HAARVI) study in Seattle, WA. Phage-DMS results from these samples were reported previously [59, 60]. This research was approved by the University of Washington Institutional Review Board (IRB number STUDY00000959).

Again, the sample set was restricted to only include individuals aged 18-55 years to better match other sample groups.

All plasma and sera were heat inactivated at 56°C for 1 hour prior to use. Full details of all samples are available in Tables 1 and S1.

### Phage-DMS, Illumina library preparation and deep sequencing

The experimental protocol was performed exactly as described previously [59]. Briefly, an oligonucleotide pool was synthesized that contains sequences coding for peptides of 31 amino acids that tile along the length of the Wuhan-Hu-1 Spike protein sequence [92] in 1 amino acid increments. For each peptide with the wildtype sequence, 19 variations were included that have a single mutation at the middle amino acid, resulting in a total library size of 24,820 unique sequences. The oligonucleotide pool was cloned into T7 phage, followed by amplification of the phage library; this step was performed twice independently to generate biological duplicate phage libraries. The phage library was incubated with a serum or plasma sample, then bound antibody-phage complexes were immunoprecipitated using Protein A and Protein G Dynabeads (Invitrogen). Bound phage were lysed, and DNA was amplified by PCR and cleaned prior to sequencing on an Illumina MiSeq or HiSeq 2500 with single end reads.

Demultiplexing and read alignment were also performed as described previously [60].

### Replicate curation

Biological replicates were analyzed in parallel to assess reproducibility of results. For simplicity, results from only one biological replicate are shown and described, with the same figures generated with the second biological replicate available to view online at https://github.com/matsengrp/phage-dms-nhp-analysis. Within each biological replicate, “in-line” technical replicates were run for some samples. In these cases, the technical replicate with the highest mapped read count was selected for analysis.

### Wildtype enrichment and defining epitope regions

The enrichment of wildtype peptides was calculated as described previously to quantify the proportion of each peptide in an antibody-selected sample relative to the proportion of that peptide in the input phage library [58]. On enrichment plots, the locus of each peptide is defined by its middle amino acid. Enrichment values of wildtype peptides were summed across epitope regions of interest for statistical comparisons between groups (“Summed WT enrichment” on figures). Mann-Whitney U tests were performed with multiple comparisons adjustment using the Bonferroni-Dunn method.

### Escape profile comparison

The effect of a mutation on antibody-peptide binding was quantified as “differential selection,” which is the log fold change in the enrichment of a mutation-containing peptide compared to the wildtype peptide. This number is multiplied by the average of the wildtype peptide enrichments at that site and its two adjacent sites to get a “scaled differential selection” value, as described previously [60]. The enrichment values of the adjacent wildtype peptides are included in this calculation to make the analysis less susceptible to noise. Negative differential selection values represent reduced binding compared to wildtype, while positive differential selection values indicate that the mutation enhanced binding. “Summed differential selection” is the sum of the 19 scaled differential selection values for all mutations at a site, and gives a sense of the overall magnitude of escape at that site.

The comparison of two escape profiles is quantified by an escape similarity score computed in the framework of an optimal transport problem [93]; this algorithm was described in detail at https://matsengrp.github.io/phippery/esc-prof.html. An overview of the method is shown in S4 Fig. Escape profiles are commonly portrayed as logo plots using scaled differential selection values (S4A Fig). At each site, escape data in logo plot form can instead be represented as binned distributions, with each mutation making some contribution to the total amount of escape at that site based on its scaled differential selection value (S4B Fig). For each site, an optimal transport problem computes the most efficient way to transform one individual’s escape distribution into that of a different individual (S4C Fig). The cost to “exchange” amino acid contributions between profiles is based on the similarity between the amino acids being exchanged, as defined by the BLOSUM62 matrix [94]. More “movement” between dissimilar amino acids drives up the total cost of the transport; therefore, a higher cost indicates less similar profiles. Escape similarity scores are the inverse of the total cost of transforming one profile into another. Scores were calculated between pairwise combinations of individuals to compare escape profile variability within and between sample groups.

### Protein structure

The structure of a SARS-CoV-2 Spike glycoprotein monomer in the closed state (PDB 6XR8) was examined to visualize epitope regions [95]. Coloring was added using UCSF ChimeraX-1.2.5, developed by the Resource for Biocomputing, Visualization, and Informatics at the University of California, San Francisco, with support from National Institutes of Health R01-GM129325 and the Office of Cyber Infrastructure and Computational Biology, National Institute of Allergy and Infectious Diseases [96].

### Code, software, and data availability

All analyses were performed in RStudio version 1.3.1093, Python version 3.6.12, GraphPad Prism version 9.0.1, and the phip-flow and phippery software suite (https://matsengrp.github.io/phippery/). The phip-flow tools perform read alignment using Bowtie2 [97] in a Nextflow [98] pipeline script. The escape profile comparisons are done with phippery in Python 3.6.12 and depend on the NumPy [99], pandas [100, 101], xarray [102], POT [103], and biopython [104] packages. All code and instructions for running this analysis are available at https://github.com/matsengrp/phage-dms-nhp-analysis.

## Acknowledgements

We thank Caitlin Stoddard for helpful guidance on the analysis and interpretation of our results. We are also grateful to Cassie Sather and others at the Genomics core facility for assistance with sequencing. We thank Lisa Jackson (Kaiser Permanente), Chris Roberts, Catherine Luke, and Rebecca Lampley [National Institute of Allergy and Infectious Diseases (NIAID), NIH] for assistance obtaining the mRNA-1273 phase 1 trial vaccine samples (NCT04283461). We thank all volunteers in the phase 1 trial, as well as all research participants and study staff of the Hospitalized or Ambulatory Adults with Respiratory Viral Infections (HAARVI) study, without whom this work would not be possible.

## Supporting information

**S1 Fig: Enrichment of wildtype peptides varies by group in newly defined epitope regions.** The locus numbers are shown on the x axis, and each individual is represented in a different color.

(A) Wildtype enrichment by group from AA 526-685, spanning the CTD-N’ and CTD-C’ epitopes.

(B) Wildtype enrichment by group from AA 777-855, spanning the pre-FP, FP, and post-FP epitopes. (C) Wildtype enrichment by group from AA 1135-1204, spanning the SH-H and HR2 epitopes.

**S2 Fig: Enrichment of wildtype peptides in baseline macaque samples compared to post-vaccination or post-infection samples.** The x axis indicates each peptide’s location along SARS-CoV-2 Spike protein, and each entry on the y axis is an individual sample. Sample groups are indicated on the left. The same macaques that contributed baseline samples also contributed post-vaccination or post-infection samples. All enrichment values over 20 are plotted as 20 to better show the lower range of the data. Above the heatmap, domains of Spike are shown with grey boxes, with the S1/S2 and S2’ cleavage sites indicated with arrows. The epitope regions defined in the current study are shown as colored boxes (from left to right: NTD in red, CTD-N’ in green, CTD-C’ in cyan, pre-FP in pink, FP in black, post-FP in orange, SH-H in purple, and HR2 in blue).

**S3 Fig: Differences in enrichment of wildtype peptides by macaque subgroups.** Wildtype enrichment values were summed for all peptides within each region of Spike that showed enrichment. Each point represents an individual macaque. No significant differences were found by Kruskal-Wallis test at a threshold of p=0.05. LION: Lipid InOrganic Nanoparticle; NLC: Nanostructured Lipid Carrier.

**S4 Fig: Use of optimal transport to quantify similarity between amino acid escape profiles.** (A) Profile 1 and 2 show example logo plots for two samples across the same region. Negative scaled differential selection values represent mutations that reduce antibody binding. Amino acids of the same color indicate similar chemistry (e.g., green = polar). (B) At each location (in this example, the boxed site in panel A), the profiles are represented as binned distributions where each bin corresponds to the contribution to escape for an amino acid substitution. (C) The optimal transport solution to transform one profile to the other is computed, where the cost to “exchange” an amino acid contribution in Profile 1 to an amino acid contribution in Profile 2 is derived from the BLOSUM62 matrix. For the purposes of the schematic, the number of dollar signs associated with each line denotes the relative cost of each move (i.e., more dollar signs = more costly = moving between amino acids that are less similar). (D) To quantify similarity between profiles, an escape similarity score is calculated as the inverse of the total cost to perform the transformation. For more details, see https://matsengrp.github.io/phippery/esc-prof.html. Created with BioRender.com.

S5 Fig. Logo plots for all vaccinated macaques, vaccinated humans, and convalescent macaques in the CTD-N’ epitope region.

**S6 Fig. Logo plots for all vaccinated humans, convalescent macaques, and convalescent humans in the FP epitope region.**

**S7 Fig. Logo plots for all vaccinated macaques, vaccinated humans, convalescent macaques, and convalescent humans in the SH-H epitope region.**

**S8 Fig. Logo plots for all convalescent macaques in the pre-FP and post-FP epitope regions.**

**S9 Fig: Differences in enrichment of wildtype peptides in vaccinated humans and convalescent macaques.** As in Fig 2, wildtype enrichment values were summed for all peptides within each epitope region of Spike. Multiple Mann-Whitney U tests were performed, with p values corrected for the number of comparisons (8) using the Bonferroni-Dunn method. ****, p ≤ 0.0001; ***, p ≤ 0.001; **, p ≤ 0.01; *, p ≤ 0.05.

